# The Spatial Scale of Synaptic Protein Allocation during Homeostatic Plasticity

**DOI:** 10.1101/2020.04.29.068833

**Authors:** Chao Sun, Andreas Nold, Tatjana Tchumatchenko, Mike Heilemann, Erin M. Schuman

## Abstract

An individual neuron hosts up to 10,000 individual synapses that can be made stronger or weaker by local and cell-wide plasticity mechanisms, both of which require protein synthesis. To address over what spatial scale a neuron allocates synaptic resources, we quantified the distribution of newly synthesized proteins after global homeostatic upscaling using metabolic labeling and single-molecule localization (DNA-PAINT). Following upscaling, we observed a global increase in locally synthesized nascent protein in synapses and at dendrites, with a high degree of variability between individual synapses. We determined the smallest spatial scale over which nascent proteins were evenly distributed and found that it is best described by synaptic neighborhoods (~ 10 microns in length)-smaller than a dendritic branch and larger than an individual synapse. Protein allocation at the level of neighborhoods thus represents a solution to the problem of protein allocation within a neuron that balances local autonomy and global homeostasis.

---

Biological compartments can function and adapt with autonomy by localizing cell biological organelles and machinery. Perhaps the best example of this is the neuronal synapse, where local control of protein production and degradation allows for the compartmentalization of synaptic function and plasticity (*1*–*4*). This creates a logistical challenge for the neuron: it must allocate its proteins between thousands of synapses to maintain their parallel computing while still fulfilling individual plasticity demands. It remains unknown how a neuron accomplishes this large-scale resource allocation to balance local autonomy (*3*–*5*) with global homeostasis (*6*). These questions are particularly important during homeostatic scaling— where cell-wide activity manipulations can result in a global multiplicative scaling of all of a neuron’s synapses (*6*). If plasticity is induced globally but proteins are distributed locally, what are the rules that govern their distribution? Understanding synaptic resource allocation, in particular the rules that govern the delivery of proteins, is crucial for understanding how local mechanisms can be implemented without compromising global homeostasis (*5*, *7*–*10*).

To address this question, we briefly metabolically labeled nascent proteins and used a single-molecule localization strategy (DNA-PAINT) to visualize and quantify their synaptic allocation in mature, cultured rat hippocampal neurons. We examined the allocation of nascent protein following a global plasticity-inducing perturbation, homeostatic synaptic upscaling (*6*), and found that new synaptic proteins were heterogeneously distributed between synapses despite a global elevation of dendritic protein synthesis. We reveal that neighborhoods of synapses, not individual synapses, comprise the relevant units for nascent-protein allocation during upscaling.

To quantify the allocation of newly synthesized proteins under control conditions and following synaptic scaling in cultured hippocampal neurons, we combined nascent protein metabolic labeling with quantitative single-molecule localization microscopy, DNA-PAINT (*11*, *12*). To tag the nascent proteome for quantitative single-molecule localization, we used a brief (15 min) pulse of the noncanonical amino acid Azidohomoalanine (AHA) (Fig. 1A; BONCAT; See Methods) (*11*). A brief labeling period was chosen because prolonged labeling created too much signal crowding, making it difficult to isolate and analyze individual clusters of nascent-protein localizations (Fig. S1). After AHA labeling, neurons were fixed and AHA-tagged nascent proteins were labelled with a single-stranded DNA “docking” oligo using Copper-free click chemistry (see Methods) (*13*). The samples were then immunolabelled for dendritic and synaptic reference markers, MAP2 and PSD-95, and fiducial markers for drift-correction following single-molecule localization analyses (*12*). Tagged nascent proteins in the dendrite were detected by repeated brief fluorescence detection events representing the transient hybridization between the docking and imager (complementary, fluorescent) oligos (Fig. 1B). As previously reported (*11*), without AHA incorporation, we observed a low level of background signal in the dendrites (Fig. 1C&D; top images indicate nascent-protein localizations; bottom images indicate the location of the dendrite), indicating that our localizations represent newly synthesized proteins. Using this method, tagged nascent proteins appeared over time as clusters of fluorescent localizations (Fig. 1E; indicated by grey arrows) in dendrites (Fig. 1E; outlined by black dashed lines). Our analyses indicate that every copy of nascent protein (shortened as ‘protein’ from hereon) is roughly equivalent to 7 ± 4 localizations (Fig. S2A; See also Methods) with an average localization precision of 13.1 nm (See Methods) and negligible bleaching effects (Fig. S2B) (*14*, *15*).

**Fig. 1.**
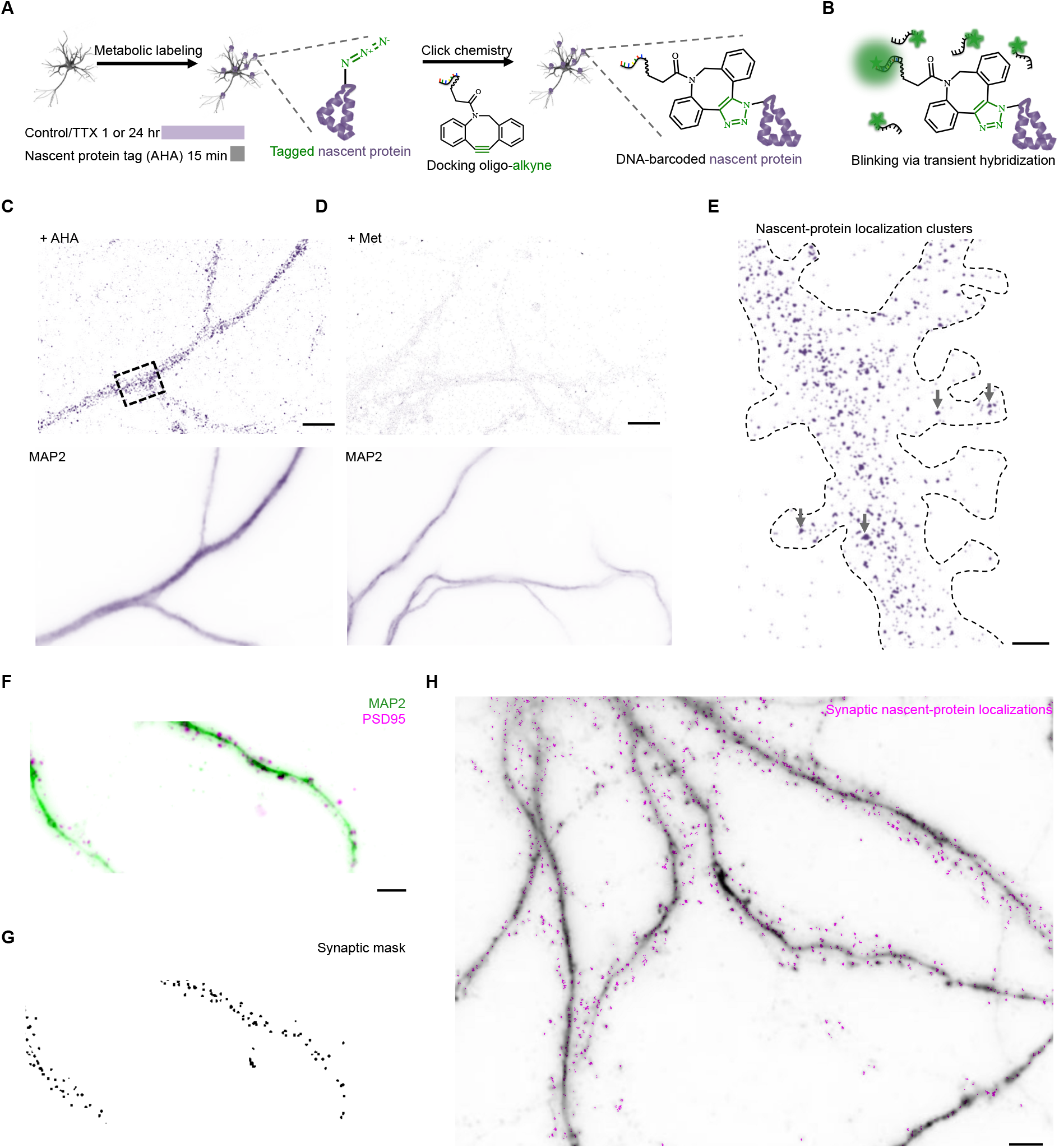
Quantitative, single-molecule localization of nascent synaptic proteins. **(A)** Cultured rat hippocampal neurons (18-21 DIV) were treated with the azide-bearing amino acid (AHA) to label nascent proteins following 1 hr or 24 hr upscaling, or mock treatment (control). Cu-free click chemistry was used to conjugate nascent proteins with a single-strand DNA oligo barcode for subsequent visualization. **(B)** Recurring, transient hybridization between the DNA-oligo barcode in A and its fluorescent complementary strand enabled single-molecule localization imaging of nascent proteins. **(C)** Representative image showing the nascent protein localization detected following treatment with AHA and the click reaction (upper image). Box indicates the region enlarged in E. Lower image shows the corresponding MAP2 immunostaining for the same region of interest. Scale bar = 6 µm. **(D)** Representative image showing the absence of nascent protein localization detected following the control treatment with Methionine instead of AHA, and the click reaction (upper image). Lower image shows the corresponding MAP2 immunostaining for the same region of interest. Scale bar = 6 µm. **(E)** Higher magnification image of dendritic segment (indicated by box in C) containing clusters of nascent-protein localizations. 4 example clusters are indicated by grey arrows; the entire dendrite segment is outlined by black dashed lines. Scale bar = 0.5 µm. **(F)** Wide-field micrograph of a neuronal dendrite after 24 hr upscaling immunolabeled with dendritic and synaptic reference markers (MAP2 in green and PSD-95 in magenta). Scale bar = 5 µm. **(G)** Example of a synaptic mask generated by a custom-written local thresholding algorithm (see Methods) for the dendritic arbor shown in F. **(H)** Synaptic nascent-protein localizations (magenta) and dendritic and synaptic reference fluorescence (gray). Scale bar = 4 µm.

Next, we quantified the extent to which individual proteins (nascent proteins) occupied excitatory synapses. The immunolabeling of dendrites (anti-MAP2; Fig. 1F) and excitatory synapses (anti-PSD-95, a postsynaptic marker; Fig. 1F) allowed us to identify and mark their position (see Methods) (*16*). Using these data, we created a synapse template, marking the position of each synapse within a dendrite (Fig. 1G). Our method detected an average synapse density of ~1 /µm (Fig. S3A), consistent with previous data (*17*). Using the synapse template, we then extracted nascent synaptic protein localizations (shortened as ‘synaptic proteins’ from hereon and used as proxy for protein copies) over a 6400 µm^2^ field–of-view (containing 10^2^-10^3^ synapses belonging to the same dendrite arbor; Fig. 1H; pink indicate synaptic proteins). Using a custom cluster identification algorithm (see Methods), we then obtained the spatial location and abundance of nascent proteins at each synapse on a dendrite (Fig. 1H) for further analyses. Protein copy numbers at synapses ranged in abundance from 0 to ~7 (0-45 localizations, see Fig. S2A and Methods) for the 1000 synapses measured from four cell replicates under control conditions. The number of nascent proteins within dendritic branches did not exhibit a gradient reflecting proximity to the cell soma (Fig. S2C; see also Fig. 1C), suggesting that most nascent proteins we detected likely arose from dendritic protein synthesis (as expected with the brief metabolic labeling used) (*2*).

To examine whether the allocation of newly synthesized proteins is changed during global plasticity induction and expression, we induced synaptic upscaling (*6*). Cultured hippocampal neurons were incubated with 2 µM tetrodotoxin (TTX) for 1 or 24 hr, representing an early “induction” time-point (1 hr) as well as a time-point when synaptic scaling had been established (24 hr) (*18*). At the end of either (1 or 24 hr) TTX incubation period, neurons were metabolically labelled (4 mM AHA) for the last 15 min before fixation and processing to identify dendrites and synapses, as described above (Fig. 1A). The synapse size we measured (approximated by PSD puncta area-see Methods) exhibited the classic log-normal distribution in both control and treated groups (Fig. S4), comparable to previously reported size ranges (*17*, *19*). As observed previously (*20*), synapse density (synapses per micron in the dendrite) increased significantly after 1 and 24 hr of activity blockade (Fig. S3A&B) with no significant change in mean synapse size (Fig. S4).

To address in which subcellular compartments nascent proteins were modified by plasticity, we quantified the amount of protein delivered to synapses and the dendrite. At synapses, we detected a significant increase in nascent protein after both 1 hr and 24 hr of upscaling (Fig. 2A and Fig. S5A-E). A significant increase in protein was also observed in whole dendrites (density per µm dendrite; Fig. 2B; also see Methods) (*21*). We evaluated the fractional allocation of proteins to different compartments: in untreated neurons we found ~5% or less of the total dendritic nascent protein present in the synapse (Fig. 2C), and the rest in the dendritic shaft. The synaptic protein fractions (Fig. 2C) increased significantly after 1 hr upscaling but not after 24 hr upscaling, despite the slight increase in synapse density in both groups (Fig. S3A). These data suggest that nascent proteins begin to congregate at and near synapses shortly after experiencing a loss of activity (Fig. 2D upscaling 1hr). By the time synaptic scaling is typically reliably detected at synapses (24 hrs) (*18*), the total dendritic nascent protein level (Fig. 2B) increased concomitantly with the protein levels at synapses (Figure 2D upscaling 24 hr); as such, the fraction of protein allocated to synapses was not significantly different from control.

**Fig. 2.**
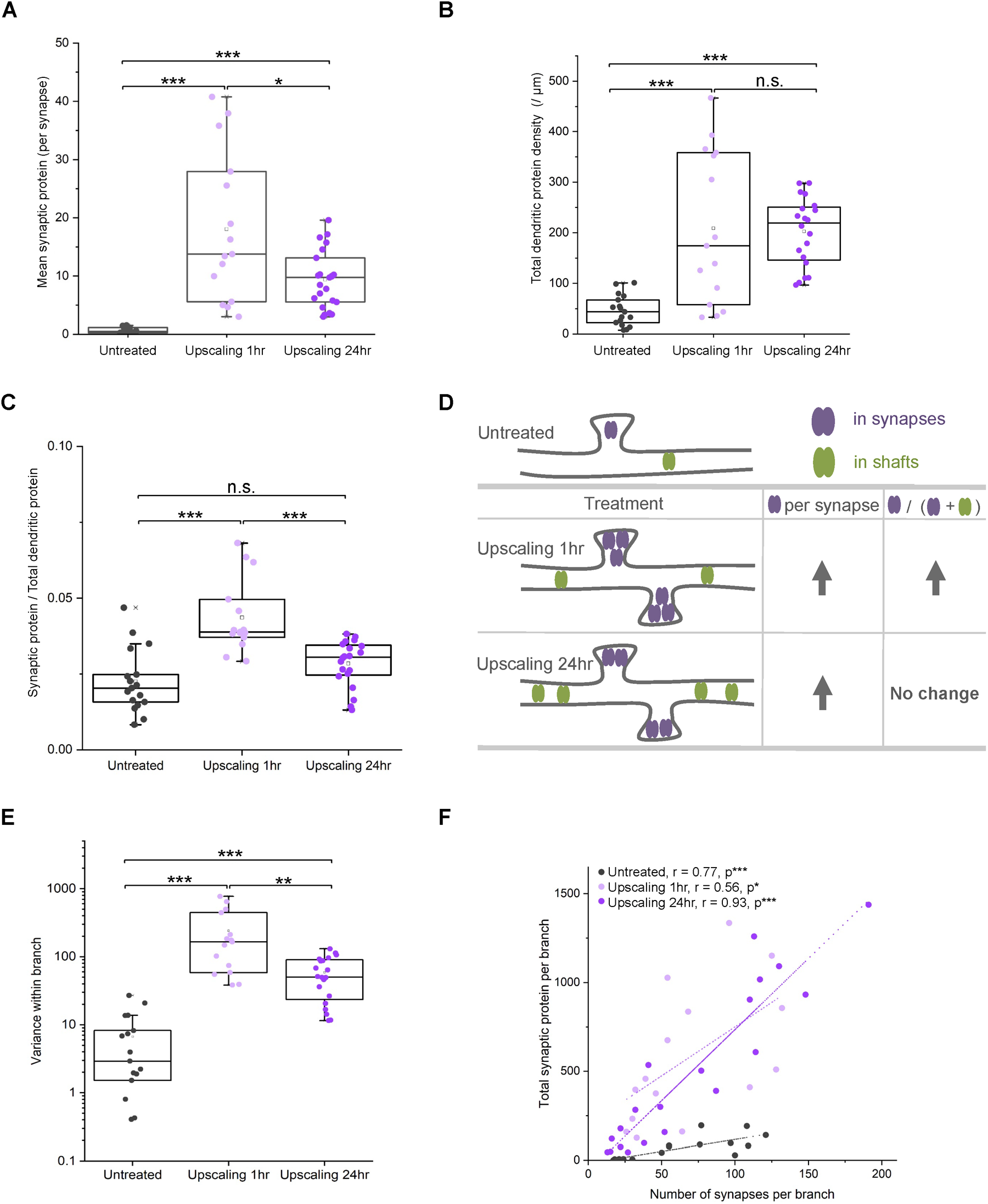
Upscaling induced an increase in nascent proteins within dendrites and at synapses. **(A)** Scatter plots indicating the mean synaptic protein (localizations; per synapse) for 17 dendritic branches from 4 untreated cells (grey), 15 branches from 3 cells treated with 2 µM TTX for 1 hr (lavender), 21 branches from 3 cells treated with 2 µM TTX for 24 hr (purple) (Same for 2B-E). Unpaired T-tests corrected for multiple comparisons (Bonferroni adjustment) indicated significant increases after both 1 hr and 24 hr upscaling (*p<0.0167; **p<0.0033; ***p<0.00033; the same applies to 2B-C & E). Box plots include the median (50^th^ percentile) indicated by the middle line in the box, the interquartile range (25^th^ to 75^th^ percentile) indicated by the top and bottom outlines of the box, the maximum and minimum without outliers (outliers defined as > 75^th^ percentile plus 1.5 folds of the inter quartile range or < 25^th^ percentile minus 1.5 folds of the interquartile range). The same description applies for all following box plots unless otherwise noted. **(B)** Scatter plots indicating the dendritic protein (localization) density (per µm dendrite). There were significant increases after both 1 hr and 24 hr upscaling (Unpaired T-test corrected for multiple comparisons). **(C)** Scatter plots showing the ratio of synaptic/total dendritic proteins (localizations). There was a significant increase after 1 hr upscaling (Unpaired T-test corrected for multiple comparisons). **(D)** Schematic summarizing the distribution of nascent protein allocation at excitatory synapses and in dendrites during upscaling. In comparison to untreated, both upscaling 1 hr and 24 hr groups exhibited an increase in synaptic nascent proteins (purple). However, the fraction of synaptic proteins only increased significantly after 1 hr upscaling but not after 24 hr due to a concomitant elevation in the dendritic proteins (green) after 24 hr upscaling. **(E)** Scatter plots indicating the variance of synaptic protein allocation within dendritic branches. There were significant increases after both 1 hr and 24 hr upscaling (Unpaired T-test corrected for multiple comparisons). **(F)** Scatter plots showing significant correlations between the synaptic protein (localizations) of each branch (y-axis) and the number of synapses (x-axis) for untreated synapses (black), 1hr upscaling synapses (lavender), and 24hr upscaling synapses (purple) with corresponding adjusted Pearson’s correlation coefficients r, and p value ranges for one-tail tests (* and *** correspond to p<0.05 and p<0.001, respectively).

While at the population level there was a scaling-induced increase in protein level at synapses and along dendrites, there was significant diversity in synaptic protein levels within the dendrite. For example, the synaptic protein within individual dendritic branches exhibited significantly higher variance following 1 hr and 24 hr upscaling (Fig. 2E; see Fig. S3C for branch definition), indicating greater differences in nascent protein allocation between synapses. To normalize for the contribution of the overall increase in nascent protein after upscaling, we calculated the coefficient of variation (CV; standard deviation normalized by mean). The CV decreased significantly following 1 hr and 24 hr upscaling (Fig. S5G), indicating that elevated synaptic protein allocation could account for the increased variance. Furthermore, the CV values remained high for the majority of dendritic branches following upscaling (above 1, namely standard deviation is greater than mean, in Fig. S5G; grey dashed line indicates CV = 1). Taken together, these results demonstrate that synaptic scaling results in a global increase of dendritic protein synthesis accompanied by a persistent and pronounced diversity in synaptic protein allocation within dendrites.

The above observed diversity, at first glance, appears at odds with the global (presumed cell-wide) nature of homeostatic scaling. Does heterogeneity exist at large spatial scales (e.g. between dendritic branches) with increasing homogeneity at smaller spatial scales (e.g. between neighboring synapses)? Or is it the opposite— global homogeneity with local heterogeneity? To answer these questions, we investigated the spatial rules for allocating proteins to synapses. At the scale of the dendrites that comprise a single neuron, we hypothesized that synaptic protein must be allocated so as to ensure the function of each dendritic branch (taking into account its length and synapse density). Indeed, the total synaptic protein of a dendritic branch was linearly correlated with its number of synapses (Fig. 2F) and its length (Fig.S5F). This suggests that synaptic protein allocation within a neuron is homogeneous across dendrites and that the relevant unit for protein allocation is at a scale smaller than an entire dendritic-branch.

At the individual-synapse scale, we considered whether all synapses within dendrites receive an equal fraction of newly synthesized protein. To understand what influences a synapse’s share of the total protein allocation, we analyzed the neighborhood factors including the local synapse density (per µm of dendrite) (*22*), and the local synaptic protein density or, in other words, the level of synaptic proteins present at neighboring synapses (per µm of dendrite). The synaptic protein density captures the mean synaptic protein allocation normalized by dendritic length and reflects a dendritic neighborhood’s propensity to acquire synaptic proteins. Due to the discrete nature of each synapse’s contribution to synapse density and protein density along the dendrite, reasonable measures of local density fluctuations are not possible without spatial smoothing. Since synapse densities in the majority of dendritic branches were higher than 1 per 2 µm (Fig. S3A), we approximated every synapse’s density contribution to a unit Gaussian with a standard deviation, σ = 2 µm (Fig S6). This spatially-limited smoothing was sufficient to measure the local synapse density (Fig. SB) without losing local features due to over-smoothing (e.g. with a higher σ value that does not match the measured synapse densities). We used a similar method to estimate every synapse’s contribution to synaptic protein densities of their neighborhoods (see Methods). Using these metrics, we analyzed each individual synapse’s share of protein in relation to its local synapse density and synaptic protein density (Fig. 3). These two local densities focus on much smaller spatial scales than branches (~2 µm around individual synapses in contrast to ~20-130 µm for branches in Fig. S5F). A 3D scatter plot depicting these three measures allows one to visualize the variability between synapses and whether a synapse’s protein allocation is determined by its local features. These plots revealed that previously described diversity in synaptic protein allocation (Fig. 2E&S5G) was also evident at the individual-synapse scale— local features such as local synapse density and synaptic protein density poorly predicted an individual synapse’s protein. Overall, the greater than 1000 synapses we measured were distributed continuously in the parameter space defined by a synapse’s protein allocation (i.e. synaptic protein localizations; z axis), local synapse density (y axis), and local synaptic protein density (x axis) (Fig. 3A-C; See also Fig. S7). During 1 hr and 24 hr upscaling (Fig. 3B&C), there was a clear increase in synaptic protein allocation (depicted by z axis) in comparison to control (Fig. 3A), consistent with Fig. 2B. We also noted that synaptic proteins after 1 hr upscaling scattered more broadly than after 24 hr (Fig. 3B; See also Fig. S7J-L normalized per dendrite), showing higher variability in protein allocation. The y-z projection illustrates the correlation between a synapse’s protein allocation level (z) and local synapse density (y); we found no correlation across treatment groups (Fig. 3D; Also see Fig. S7D-F & M-O), indicating that local synapse density does not predict how much protein a synapse obtains. Finally, the x-z projection illustrates the correlation between a synapse’s protein allocation (z) and its neighboring synapses’ propensity to acquire proteins (x); we found only weak correlations here (Fig. 3E; also see Fig. S7G-I & P-R) with or without upscaling. Such diversity of synaptic protein allocation among neighboring synapses is reminiscent of the competition among neighboring synapses previously reported (*23*). These observations were robust across dendrites and neurons within the same treatment group (Fig. S7). Overall, these data indicate that protein allocation is locally heterogeneous— individual synapses exhibited a high level of autonomy. If so, how is global homeostasis achieved given this observed local heterogeneity?

**Fig. 3.**
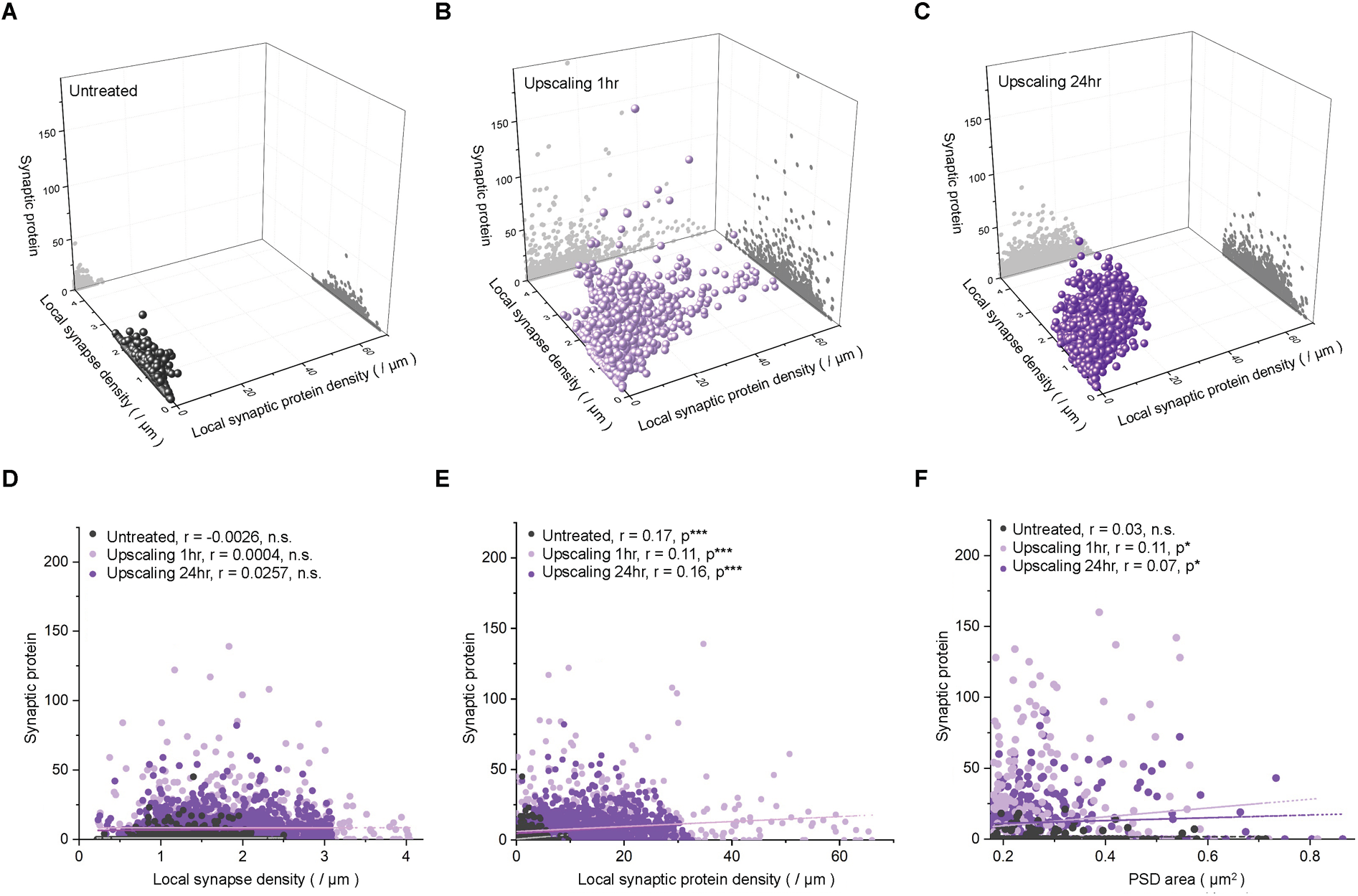
Synaptic protein allocation poorly correlated with local synapse density, local synaptic protein localization density, and synapse size. **(A)-(C)** 3D scatter plots of **(A)** 1000 synapses from four untreated neurons, **(B)** 1037 synapses from three 1 hr upscaling neurons, and **(C)** 1414 synapses from three 24 hr upscaling neurons. x-axis: local synaptic protein density from neighboring synapses (see Methods and Fig. S6); y-axis: local synapse density (also see Methods and Fig. S6); z-axis: synaptic protein (localizations) of a synapse. Projections on x-z and y-z planes are colored in dark and light grey, respectively. **(D)** Scatter plots showing the absence of significant correlations between synaptic protein (localizations) of each synapse (y) and the measured local synapse densities (x) for untreated synapses (black), 1 hr upscaling synapses (lavender), and 24 hr upscaling synapses (purple) with corresponding adjusted Pearson’s r values and p value ranges for one-tail tests. **(E)** Scatter plots indicating weak correlations between synaptic protein (localizations) for each individual synapse (y-axis) and the measured local synaptic protein (x-axis) for untreated synapses (black), 1 hr upscaling synapses (lavender), and 24 hr upscaling synapses (purple) with corresponding adjusted Pearson’s r values and p value ranges for one-tail tests (*** corresponds to p<0.001). **(F)** Scatter plots showing weak correlations between synaptic protein (localizations) for each individual synapse and its corresponding measured PSD puncta sizes in control (black, 225 puncta), upscaling 1 hr (lavender, 555 puncta), upscaling 24 hr (purple, 428 puncta) with corresponding adjusted Pearson’s r values and p value ranges for one-tail tests (* corresponds to p<0.05).

We hypothesized that the existing landscape of synapse weights may dictate synaptic need and, consequently, maintain global homeostasis during upscaling (*6*). Therefore, using synapse size as a proxy for synaptic weight (*16*, *24*), we next examined whether a synapse’s size influences its share of nascent proteins. We measured the area of the immuno-labelled PSD-95 puncta as a proxy for synapse area (*16*, *25*) (see Methods) and observed a weak correlation between the synaptic protein allocation and the measured PSD puncta size across all treatment groups (Fig. 3F). The observed weak correlation was likely due to a large population of small-to-medium-sized synapses (e.g. 0.2-0.3 µm^2^ in Fig. 3F) that exhibited a wide range of protein allocation levels (Fig 3F). Indeed, when small synapses (<0.2 µm^2^) were compared with larger synapses (>0.3 µm^2^), we detected a significantly higher protein allocation among >0.3 µm^2^ synapses in both control and treatment groups (Fig. S8), consistent with the idea that large synapses have higher protein needs. Even so, the overall small correlations in Fig. 3F suggest an uncoupling between a snapshot measurement of a synapse size (e.g. using PSD-95 immuno-labeling as a proxy) and the current or future functional efficacy of the synapse (*26*, *27*).

If individual synapses exhibit a high level of autonomy in protein allocation (Fig. 3D-F), then what maintains the global homeostatic allocation of protein observed at the dendrite-level (Fig. 2F&S5F)? Here we re-examined the spatial heterogeneity of the synapse distribution and the synaptic protein distribution along dendrites (Fig. 4A for synaptic protein density; See also Fig. S9 for synapse density; see Methods and Fig. S6). The heat-map of synaptic protein density (Fig. 4A right inset) clearly indicates heterogeneity within dendritic branches represented by local domains of high and low protein densities. Such local domains span ~ 5-10 µm, clearly larger than individual synapses (*3*, *4*, *28*). This suggests that the relevant spatial scale for synaptic protein allocation might comprise neighborhoods of synapses within dendrites (Fig. S9A. See also Fig. S9B for illustration of different spatial scales) (*4*, *29*).

**Fig. 4.**
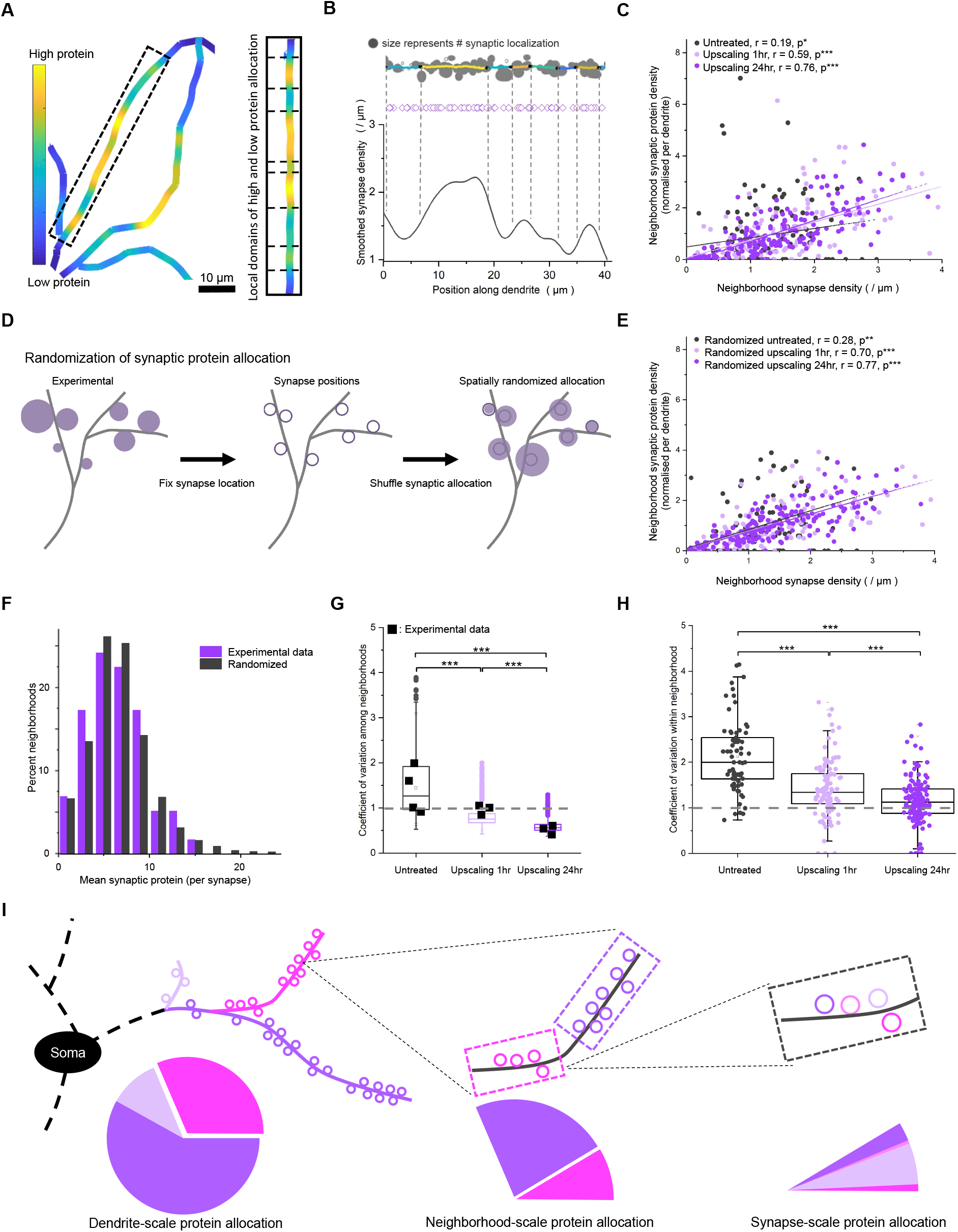
Nascent proteins were fairly allocated to synapse neighborhoods. **(A)** Heatmap of synaptic nascent-protein density (per µm dendrite; cell soma located in the bottom-right direction of the image) in a representative dendrite exhibited spatial heterogeneity at an intermediate length-scale (10^0^ - 10^1^ µm). Right inset box illustrates the emergence of local domains with high and low densities of synaptic proteins. **(B)** Synapse neighborhoods were divided based on the heterogeneity of synapse density (y-axis) along dendrites (x-axis). The peaks and valleys of the density fluctuation are identified. Borders (dashed vertical lines) between synapse neighborhoods are drawn between the peaks and valleys. The resulting division of neighborhoods matches the visible changes in synapse density (top cartoon; hollow circles represent synapses and the filled size represents protein allocation level (localization counts) in a synapse; dendritic shafts are colored and segmented by neighborhood boundaries depicted by black dots). Pink hollow spades below represent the positions of synapses along the dendritic length. **(C)** Correlations between normalized neighborhood synaptic protein (localization) density and synapse density (/ µm) in 124 neighborhoods from 4 untreated neurons (black), 130 neighborhoods from 3 upscaling 1 hr neurons (lavender), and 168 neighborhoods from 3 upscaling 24 hr neurons (purple). r values represent the Pearson’s correlation coefficient in each treatment group. p value ranges were from one-tail tests (* and *** correspond to p<0.05 and p<0.001, respectively). Every neighborhood synaptic protein density was normalized against its corresponding mean synaptic protein density (/ µm dendrite) in dendrites. **(D)** Scheme depicting the random redistribution of existing synaptic protein allocations to synapse locations within a dendrite. The size of each filled purple circle represents the amount of protein allocation. Their sizes were kept the same during randomization. **(E)** Correlations between normalized neighborhood synaptic protein (localization) density and synapse density in spatially randomized dendrites (D) in untreated (black), 1 hr upscaling (lavender), and 24 hr upscaling (purple) groups. r values represent Pearson’s correlation coefficient for each treatment group. p value ranges were from one-tail tests (** and *** correspond to p<0.01 and p<0.001, respectively). Every neighborhood synaptic protein density was normalized against its corresponding mean synaptic protein density (/ µm dendrite) in dendrites. **(F)** Frequency distributions of mean synaptic protein (per synapse) among the neighborhoods of an example dendrite from 24 hr upscaling (experimental; purple) and its 500 randomizations (black). **(G)** Box plots showing the range of coefficients-of-variation in 500 randomizations for all the neighborhoods in control (untreated; black), 1hr upscaling (lavender), and 24 hr upscaling (purple). There were significant differences between all treatment groups (Unpaired T-test corrected for multiple comparison; *** corresponds to p<0.00033). Filled black squares indicate the coefficient-of-variation among neighborhoods of the original dendrites in each treatment group. Grey dashed line indicates CV = 1. **(H)** Scatter plots showing the coefficients of variation among synapses within each neighborhood in the original data. There were significant differences between all treatment groups (Unpaired T-test corrected for multiple comparison; *** corresponds to p<0.00033). Grey dashed line indicates CV = 1. **(I)** Scheme depicting that synaptic protein allocation is globally fair at dendrite-level (left; bottom left pie chat indicates fair protein allocation among branches of corresponding colors); this is because each branch consists of multiple allocation units, i.e. neighborhoods of synapses (center; bottom center graph indicates the corresponding wedge for the magenta branch in bottom left further divided fairly among the two neighborhoods of corresponding colors); however within each neighborhood, synaptic protein allocation is heterogeneous between neighboring synapses (right; bottom right graph indicates the corresponding wedge for the magenta neighborhood in bottom center further divided unevenly among neighboring synapses of corresponding colors).

To investigate how synaptic neighborhoods acquire their share of proteins during global plasticity, we identified turning points in the synapse density fluctuations along the dendrites (Fig. S6) and segregated each dendrite into spatial neighborhoods of synapses (Fig. 4B). The high synapse-density neighborhoods (e.g. Fig. 4B yellow neighborhoods) included synapse clusters similar to what has been previously described (*4*, *7*, *28*, *30*). We measured the synapse density and synaptic protein density of every neighborhood, in addition to other characteristics such as neighborhood size, synapse number, and synaptic-protein per-synapse (Fig. S10A-C). The majority of neighborhoods were less than 10 µm in length (Fig. S10A), most of them containing < 10 synapses (Fig. S10B). Following 1 hr upscaling, there was a decrease in neighborhood size (Fig. S10D). As mentioned previously, the neighborhood synapse density describes potential loci that need proteins and therefore, can be used as a proxy for neighborhood demand. On the other hand, the neighborhood synaptic-protein density reflects a neighborhood’s ability to acquire proteins. For example, a long dendritic segment with few synaptic proteins (due to either low synapse number or low average synaptic protein allocation) would result in a low protein allocation for this neighborhood. To reveal the variability among neighborhoods rather than among the dendrites, neighborhood synaptic-protein density was normalized per dendrite. Following both 1 and 24 upscaling, we observed progressively enhanced correlations between neighborhood protein allocation (synaptic protein density) and neighborhood demand (synapse density) among 10^2^ neighborhoods (Fig. 4C; correlation coefficient r = 0.76 after upscaling 24 hr), when compared to control. Furthermore, following upscaling, the variability (CV) of neighborhood protein allocation (measured by protein-per-synapse) was reduced (Table S1). Taken together, the above indicate that the neighborhood protein allocation was likely largely determined by neighborhood synapse-number.

The above analysis suggests that protein was fairly allocated across neighborhoods under both control conditions and after homeostatic plasticity. To validate this, we randomized the coupling between a synapse’s protein allocation and its location (Fig. 4D). If a neighborhood exhibited the propensity to locally accumulate low- or high-protein allocation synapses, this randomization would eliminate this local effect. And if neighborhood protein allocation is based on synapse number only, then the experimental observations should not differ significantly from the randomized cases. Indeed, we found that after randomization, both the correlation coefficient r and the progressive increase of r over the course of upscaling were qualitatively preserved (Fig. 4E). The randomized untreated group still exhibited a lower r value (0.28) while randomized 24 hr upscaling exhibited an elevated r (0.77), similar to the experimental data (r = 0.76; Fig. 4C). We then compared the experimental distribution of neighborhood synaptic-protein per-synapse with 500 randomizations (Fig. 4F for an example dendrite and its average randomized distribution; Also See Fig. S11 for all dendrites). The box plots in Fig. 4G show the range of CV for 500 randomized distributions. We found that none of our experimental data (as dendrites; squares in Fig. 4G) was a statistical outlier. This indicates that indeed there was no significant clustering of high- or low-protein allocation synapses within specific neighborhoods, reminiscent of the lack of local homogeneity at the synapse-scale (Fig. 3D-E). We conclude that following upscaling, neighborhood protein allocation was dominated by synapse number. This further indicates that the proximity-to-the-soma did not systematically bias protein allocation, as might be expected if we sampled the proteins produced locally (see also Fig. 4A, S2C & S6D). Finally, within most neighborhoods in control and treatment groups, the CV among synapses remained high (Fig. 4H, CV =1 indicated by grey dashed line; compared with Fig. S5G), indicating heterogeneity between synapses within neighborhoods. Taken together, these data indicate that during synaptic upscaling, dendritic neighborhoods of synapses, rather than individual synapses, are a relevant allocation unit for nascent-protein distribution. Neurons use synaptic neighborhoods to manage synaptic resource allocation presumably because this strategy can accommodate diversity within neighborhoods and maintain homeostasis between neighborhoods and consequently, between dendritic branches (Fig. 4I).

Elevated protein synthesis in dendrites and synapses has been observed following several forms of plasticity (*1*, *2*, *29*, *31*). Indeed, homeostatic scaling requires new protein synthesis and a remodeling of the entire nascent proteome has been observed (*32*). We tracked the nascent protein pool following homeostatic scaling at different spatial scales: the dendrite scale (~50 µm), the synaptic-neighborhood scale (~5-10 µm), and the individual-synapse scale (~2 µm). We found an increase in locally synthesized nascent protein within synapses and dendrites after synaptic upscaling. While protein allocation at the dendritic-branch scale could be accounted for by the number of synapses and branch length, there was significant variability in the nascent protein observed at individual synapses— no strong correlations were observed with either the local synapse density or the amount of nascent protein present in neighboring synapses. As such, neighboring synapses sharing the same local environment can differ significantly in their protein allocation— each synapse determines its own protein allocation autonomously. In this regard, it is noteworthy that even synapse size failed to reliably predict synaptic protein allocation. Molecular signals that are not directly related to synapse size may be important for the capture and sequestration of nascent proteins, consistent with the idea of synaptic tagging (*23*, *33*). Such mechanisms may also decouple neighboring synapses from each other despite their common neighborhood.

The smallest scale at which nascent protein allocation was evenly distributed was small synaptic neighborhoods within dendritic branches. Neighborhoods as the relevant allocation unit imply that they have a more stable ‘identity’ than individual synapses (e.g. defined by the stable presence of diverse organelles or the sum synaptic weights). In the face of synapses’ individual variability and uncertainty (*26*), neighborhoods may represent a good scale to implement and maintain homeostasis (*4*, *7*). For example, local-scale studies of clustered plasticity in the past decade have revealed regulation within neighborhoods of synapses; long-term potentiation (LTP)-induced spine growth is accompanied by stalling and shrinkage of neighboring spines and has been proposed to balance local synaptic strengths during plasticity (*7*, *8*, *30*). This is likely accomplished by an underlying inequality in cell biological resources between neighboring synapses (*4*, *23*). Indeed, the net amount of nascent protein to potentiate and depress synapses could be balanced for a neighborhood despite the diverse demands from individual synapses or the competition between neighbors. As such, the global homeostasis in protein allocation is maintained locally by synaptic neighborhoods (*10*).

The fairness of the neighborhood allocation is consistent with local sourcing of protein derived from local ribosomes (*34*) and organelles such as mitochondria (*29*). Consequently, the spatial scale of protein allocation-units (i.e. synapse neighborhoods) might reflect the spatial scale of intracellular organelles. The observed plasticity-induced decrease in neighborhood size (e.g. following 1 hr upscaling) implies that these organelles have become more ‘local’ (e.g. redistributed closer to synapses) (*3*, *34*). Furthermore, the synapse neighborhoods exhibited similar lengths and synapse numbers as previously reported functional dendritic segments (*28*). Future studies should examine the relationship between the function of dendritic segments and synapse neighborhoods. For example, it would be interesting to see if a neighborhood’s availability of cell biological resources affects its synapses’ capacity for plasticity.

## Acknowledgments

We thank G. Tushev and L. Anneser for assistance in developing the synapse identification algorithm, A. R. Dörrbaum and S. tom Dieck for instructions during metabolic labeling and synaptic upscaling, C. Böger and A.S. Hafner for advice on image acquisition and analysis of DNA PAINT, I. Bartnik, N. Fuerst, and D. Vogel for the preparation of cultured neurons, and S. tom Dieck, M. van Oostrum, P. Donlin-Asp, J. Perez, and G. Laurent for inputs during manuscript preparation.

## Author contributions

C.S. and E.M.S. designed the experiments. C.S. conducted the experiments and image analysis. A.N. and C.S. performed the data analysis. All authors discussed the data. C.S. and E.M.S. wrote the manuscript. All authors edited the manuscript

## Data and materials availability

All data is available in the main text or the supplementary materials. Analysis scripts are available upon request.

## Supplementary Materials

Materials and Methods

Figures S1-S11

Tables S1

