## Supplemental Information for "The Spatial Scale of Synaptic Protein Allocation during Homeostatic Plasticity"

**This PDF file includes:**

Materials and Methods
Figs. S1 to S11
Tables S1

### Materials and Methods

Cell Culture

Dissociated rat hippocampal neuron cultures were prepared and maintained as described before (*21*). Briefly, rat hippocampi were dissected from postnatal day 0-1 pups of either sex (Sprague-Dawley strain; Charles River Laboratories), first dissociated with papain (Sigma) and then plated at a density of 30 $\times 10^3$ cells/cm$^2$ on poly-D-lysine coated glass-bottom Petri dishes (MatTek). Neurons were maintained (until day of experiment) in a humidified atmosphere at 37 °C and 5% $CO_2$ in growth medium (Neurobasal-A supplemented with B27 and GlutaMAX-I, life technologies) for 18-21 DIV to ensure synapse maturation. All experiments complied with national animal care guidelines, the guidelines issued by the Max Planck Society and were approved by local authorities.

Synaptic Upscaling and Metabolic Labeling with AHA

Neurons (18-21 DIV) on MatTek dishes were incubated in the growth medium described above containing 2 μM (Tetrodotoxin) TTX for 1 or 24 hrs at 37 °C and 5% CO2. 15 mins before each treatment ended, neurons were incubated in methionine-free Neurobasal-A (custom-made by Life Technologies) supplemented with 4 mM AHA (prepared as described in Link et al., 2007) and 2 μM TTX. In methionine control experiments, AHA was replaced by 4 mM methionine (Sigma). Subsequently cells were washed quickly twice with Neurobasal-A and fixed in PFA-sucrose (4 % paraformaldehyde; Alfa Aeser), 4 % sucrose in PBS-MC) at room temperature for 20 min, washed, permeabilized with 0.5 % Triton X-100 in 1 x PBS pH 7.4 for 15 min and blocked with blocking buffer (4 % goat serum in 1 x PBS) for 1 h. To optimize conditions for the subsequent click reaction, neurons were equilibrated by washes with 1 x PBS pH 7.8.

The BONCAT part of the assay was performed as described previously (*21*) with the following modification: we used a previously reported single-stranded DNA oligo sequence (P1 docking oligo) (*12*) modified to carry a more reactive alkyne, dibenzocyclooctyne (DBCO; GeneLink), as a tag in the copper-free azide–alkyne cycloaddition click reaction (*13*). For the Cu-free click reaction, 0.4 μM P1-DBCO tag was prepared in 1 x PBS (pH 7.8) before application to the cells and the click chemistry was performed overnight at room temperature. After the click reaction, cells were washed 2 times with PBS pH 7.8, 0.5 % Triton in PBS, and 3 times with PBS pH 7.4 before immunofluorescence labeling with various markers (see below). We estimated the under-sampling rate of our metabolic labeling based on the measured synaptic nascent-protein localizations (see below Data Analysis).

Immunofluorescence labeling

Neurons were washed three times in PBS before being blocked in PBS containing 4% goat serum (Gibco) for 1 hr. Neurons were incubated overnight with guinea pig antibodies anti-MAP2 (1:2000, 188004, Synaptic Systems) and anti-PSD95 (1:1000, MA1-046, Thermo Fisher) in PBS containing 4% goat serum (Gibco) at 4°C to stain the dendrites for morphology and excitatory synapses, respectively. The samples were then washed three times in PBS (5min each) before incubation for 1 h with anti-guinea pig antibody conjugated with AlexaFluor-488 (1:1000, Nanoprobes) and anti-mouse antibody with AlexaFluor-546 (1:1000 Nanoprobes). Neurons were washed three times in PBS (5 min each). All steps were performed at room temperature. Neurons were then stored in PBS at 4°C for up to three weeks until DNA-PAINT imaging.

Super-resolution microscopy

For DNA-PAINT imaging, the imaging buffer contains 500 pM P1-Atto655 (CTAGATGTAT-Atto655, Eurofins Genomics) in 500 mM NaCl, pH 7.3 (*15*).

Immunolabeled cultured neurons containing 90 nm gold fiducial markers (A1190, Nanoparz) were imaged on an N-STORM system (Nikon, Japan): an Eclipse Ti-E inverted microscope, equipped with a Perfect Focus System (Ti-PSF) and a motorized x-y stage. Total internal reflection fluorescence (TIRF) and highly inclined and laminated optical sheet (HILO) (*15*) configurations were adjusted using a motorized TIRF illuminator in combination with a 100 x oil-immersion objective (CFL Apo TIRF, NA 1.49) with a final pixel size of 158 nm. For imaging, 647 nm excitation wavelength was used, housed in a MLC400B (Agilent) laser combiner. An optical fiber guided the laser beam to the microscope body and via a dichroic mirror (T660LPXR, Chroma) to the sample plane. Fluorescence emission was separated from excitation light via a bandpass filter (ET705/72m, Chroma) and detected by an iXon Ultra EMCCD camera (DU - 897U-CS0-23 #BV, Andor). The software NIS-Elements Ar/C and µManager were used to control the setup and the camera (*15*).

Wide-field micrographs of the MAP2 and PSD95 reference markers (see above Immunofluorescence labeling) were obtained before DNA-PAINT as summed projections of 2-µm thick Z-stacks to capture the entire dendritic volume. HILO illumination was used for super-resolution acquisitions with a power of 30-40 mW, which was determined directly after the objective and under wide-field configuration. The intensity density (45% intensity of 647 nm laser) was 0.9 kW/cm2. Time-lapse datasets with 50,000 frames and 16 bit depth were acquired at 5 Hz frame rate and 5 MHz camera read-out bandwidth; pre-amplification: 3; electron multiplying gain: 4.

Data Analysis

DNA-PAINT acquisitions were reconstructed with Picasso:Localize, a module of the Picasso software (*12*), by applying a minimal net gradient of 2500. With Picasso:Render, drift corrections were applied in 2 subsequent fashions: First, a drift correction based on the redundant cross-correlation (RCC), with a segmentation of 200 was applied. Secondly, fiducial markers (gold beads) were manually selected, localized, and used for drift correction. Drift-corrected data was filtered using Picasso:Filter. Raw localizations within the same location were further filtered temporally based on their average frame-numbers to eliminate background signal due to the unspecific binding of the imger oligo to a random target. Such background signals are often clustered temporally rather than distributed through the imaging course, resulting in lower or higher average frame-numbers and lower frame-number variance compared to real signal. Afterwards, raw localizations within a maximal distance of 6X measured localization precision and showing a maximum number of transient dark frames of 20 were linked together, resulting in a single, linked localization event (referred to as ‘localization’). Localization precision (NeNA values) was determined to be 13.1 nm, as previously described using nearest-neighbor based analyses (*14*).

Synaptic regions were determined using a custom-written algorithm and the diffraction-limited PSD95 immunolabeling signal. To identify the positions of excitatory synapses, local PSD95 puncta maxima and minima were identified and normalized to the same intensity range (0 for minima, and 255 for maxima). Pixels of PSD95 puncta with over 12% intensity (30 in normalized intensity) of its associated, normalized local maximum (255 in normalized intensity) were selected to define the synaptic compartments. Such local intensity thresholds resulted in puncta size estimates that were less affected by local intensity and background differences (e.g. the phenomena where brighter puncta appear larger by eye while less bright puncta appear smaller; Fig. S4 right table) due to heterogeneity in staining or focusing. Puncta were eliminated if on the soma or more than 2 μm away from a dendritic shaft marked by MAP2 signal; Puncta with sizes smaller than 0.15 μm2 were excluded from size-based analyses due to significant inaccuracy of size measurements at smaller spatial scales. Dendrites with high level of background signal outside MAP2-labeled regions, dendrites containing overlapping signals with AHA labeling from glia, out-of-focus dendritic branches, and branches shorter than 5 μm were all eliminated. All in all, this created a synapse mask with a measured, average synapse density and synapse size distribution consistent with published values (*17*). This mask effectively enriched protein signals allocated into the synaptic area by excluding adjacent regions such as nearby proteins in the shaft (e.g. spine base). To evaluate adjacent regions such as the spine neck and base, a mask was generated to include an area of 1 μm in diameter centered on the PSD intensity maxima (Fig. S5A; subtracting the associated synaptic area). All localizations < 2.5 μm from a skeletonized line that traversed the center of the MAP2-labeled dendrites were included as total dendritic localizations (Fig. S5A).

Synaptic nascent-protein localizations were further identified by a custom-written, noise-tolerant cluster identification algorithms (DBSCAN-based) (*35*), the spatial-distance parameter (r) of which was determined based on the measured localization precisions for each dendrite. In particular, any localization with < 3 localizations within r was excluded from clustering, which is more selective than DBSCAN (*35*). It is therefore more robust against the inclusion of noise localization (see A.N., C.S., E.M.S., M.H., and T.T. in preparation for an extensive validation and comparisons with existing methods). The PAINT signal of a single docking oligo (further approximated as a single protein copy due to the low incorporation efficiency of AHA) was determined as such: We plotted the number-of-linked-localization distributions (see Fig. S2) for all the identified synaptic clusters and attributed the smallest population of clusters as the signal of a single protein copy; we further determined the average dark time for a single protein copy from this population (containing <=11 linked localizations) (Fig. S2) and found it comparable with previously reported benchmark test of a single docking oligo based on DNA origami (with identical set-up) (*15*); as such, we calculated that a single protein copy corresponds to roughly 7 ± 4 linked localizations. Due to these approximations and the known difficulties in obtaining accurate molecule counting (*36*), in the main text, we directly use the number of linked localizations as a proxy measure for protein copy numbers. In parallel, the widely used DBSCAN cluster identification (*35*) was also implemented to double-check the robustness of our subsequent analyses. Afterwards, Picasso:Render and ImageJ were used for dendritic-branch level analyses. A one-dimensional dendritic tree was constructed based on the MAP2-immunolabeling signal. Synapses and localization positions were then mapped onto the closest dendritic branch. The local synapse and synaptic nascent-protein densities per unit length were obtained by applying smoothing with a Gaussian kernel with parameter sigma = 2 μm. Note that for protein density, this contribution is proportional to its synaptic localization counts. To define neighborhoods, borders were drawn at the turning-points of the smoothed synapse density, such that neighborhoods with a range of low and high synapse densities were identified. We constrain neighborhood size to be at least 0.5 μm, and the slope of the synapse density at the turning points to be > 0.1 synapse per μm2 for partitioning into a new neighborhood. The synapse densities and synaptic nascent-protein densities were measured using the corresponding total numbers divided by the neighborhood dendritic lengths. To estimate the under-sampling rates due to metabolic labeling and click chemistry, we note that in untreated groups, synaptic nascent-protein localization was ~ 1-2 per synapse. A synapse contains approximately $10^4$-$10^5$ protein copies with an average half-life of 5 days in vitro (*37*). This gives an estimate of 1-10 proteins being renewed every 15 min, which amounts to ~ 7-70 linked localizations per synapse (Fig. S2; assuming ~ 1 localization per synapse in untreated) (*15*). This yields an under-sampling estimate of 86%-98.6%, meaning that we likely acquire data from 1.4%-14% of the synaptic nascent protein pool. Under-sampling is crucial for avoiding high density, overlapping localizations, which may introduce inaccuracies in quantification (See Fig. S1).

Reagents

Unless noted otherwise, all substances were molecular biology or cell culture grade and purchased from Sigma-Aldrich or Roth. TTX citrate was used from stock solutions (Tocris, 2 mM in H2O, 2 µM final).

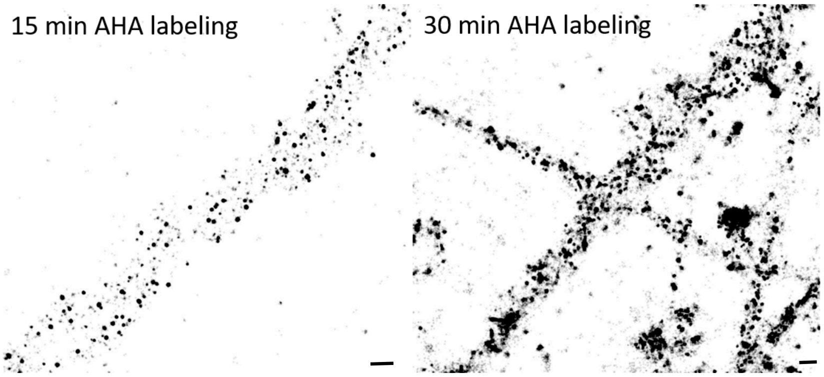


**Fig. S1.**

**DNA-PAINT images showing localizations of nascent proteins after different AHA labeling periods.** Left: after a 15 min AHA labeling, as used in this paper. Right: after a 30 min AHA labeling showing crowding and merging of signal clusters, compromising quantitative analyses. Scale bars: 0.5 µm.

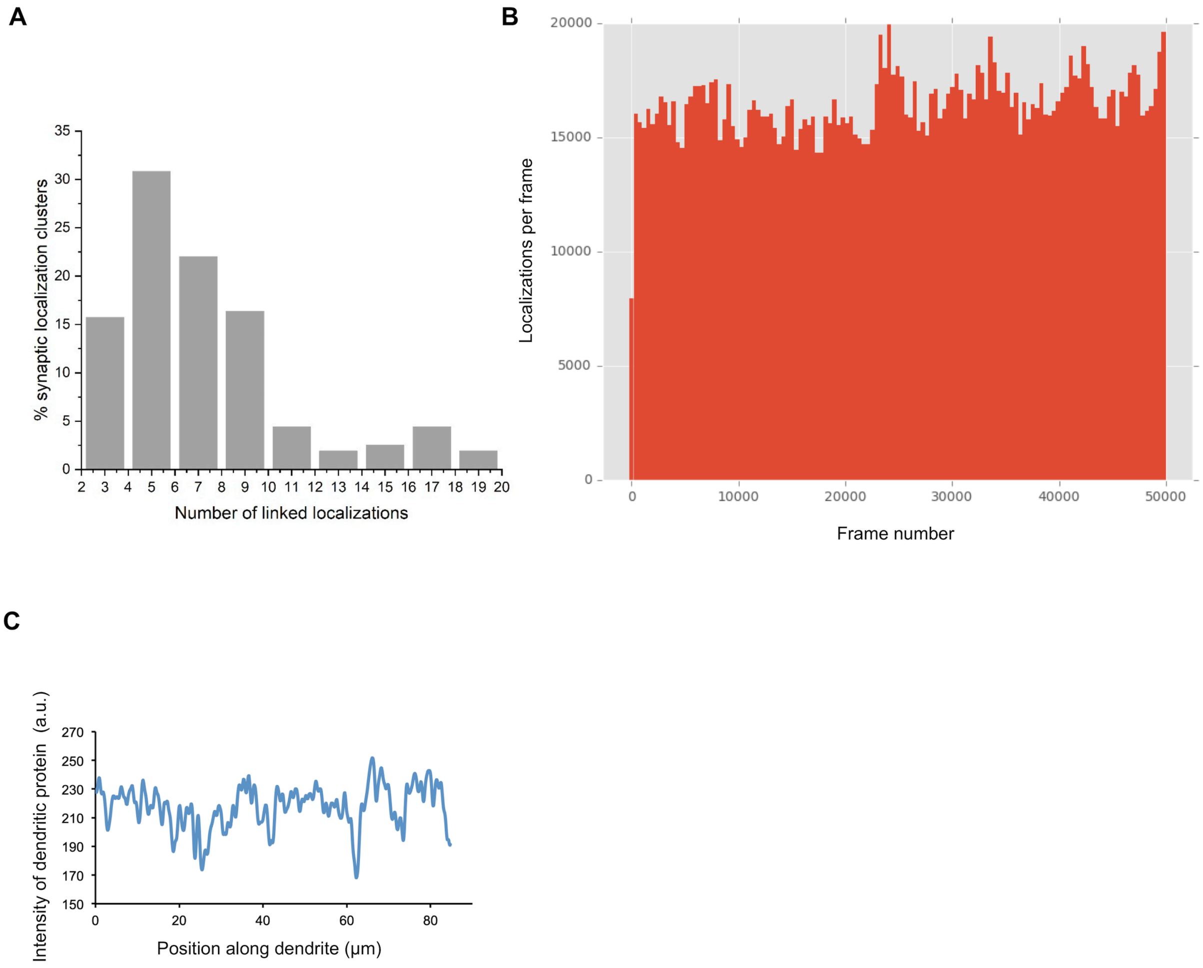


**Fig. S2.**

**Characterization of DNA PAINT clusters. (A)** Distribution of localization clusters at individual synapses. We attributed the smallest population of clusters (bins between 3 and 11 linked localizations; bin size: 2) as one protein copy carrying a single docking oligo. The average dark time ($\tau_{dark,\ single}$ ~ 1349 s) estimated for a single docking oligo is comparable with the average dark time measured previously for a single docking oligo under similar imaging conditions (ref No. 15 in main text; also see method). The larger-cluster population (e.g. 16-18 localization bins) likely contains more than one docking oligo due to multiple copies of proteins (see methods). **(B)** Localization counts per frame showed a negligible decrease over 50,000 frames, indicating negligible bleaching. **(C)** Line graph showing the local fluctuations in dendritic protein level approx. by the super-resolved image intensity (arbitrary units, a.u.) along a secondary dendritic branch (5-µm thick line profile).

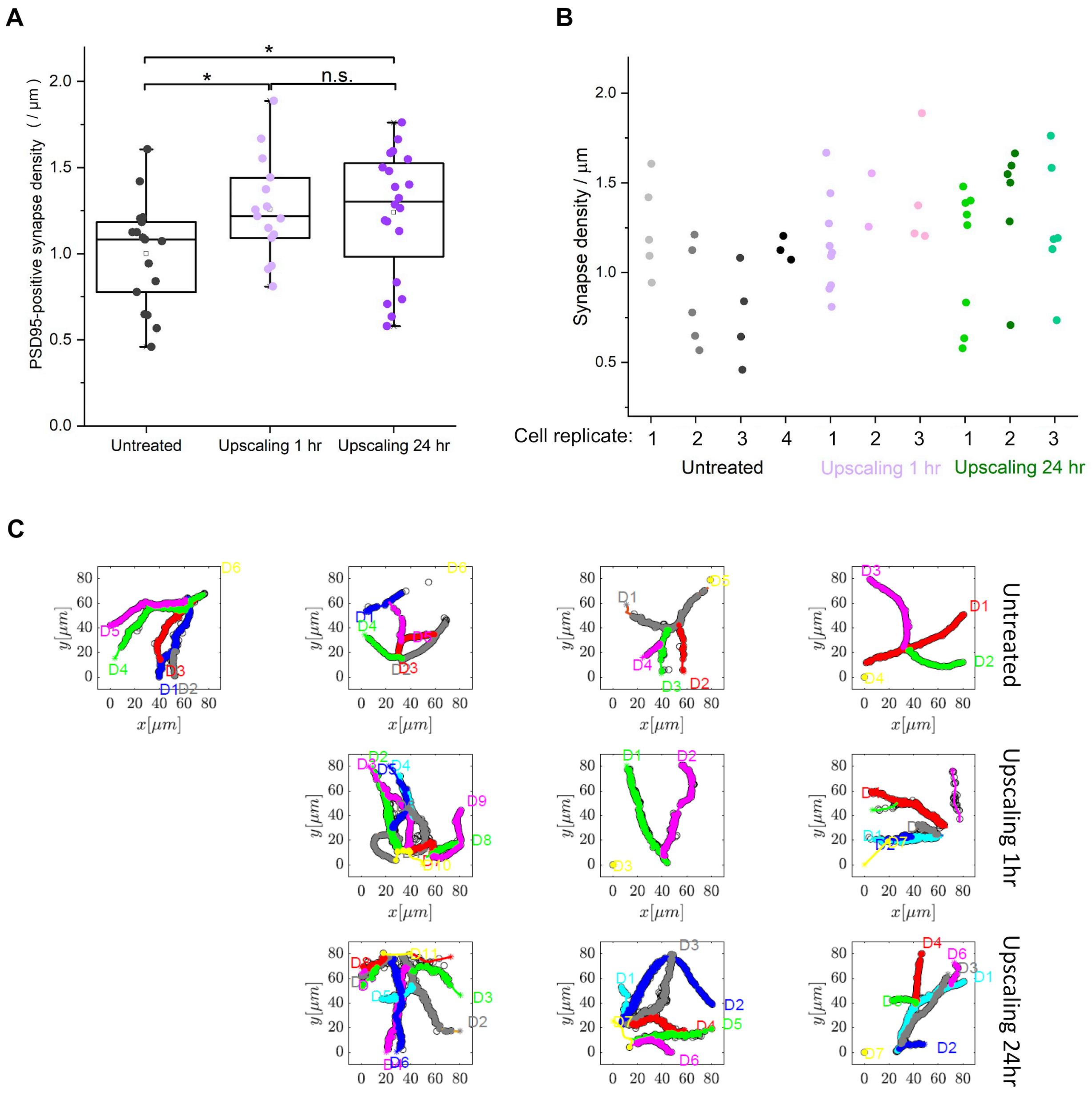


**Fig. S3.**

**Synapse density of all dendritic branches. (A)** Scatter plots indicating the synapse densities (per µm dendritic length) for 17 dendritic branches from 4 untreated cells (grey), 15 branches from 3 cells treated with 2 µM TTX for 1 hr (lavender), 21 branches from 3 cells treated with 2 µM TTX for 24 hr (purple). Unpaired T-tests corrected for multiple comparisons (Bonferroni adjustment) indicated a significant increase in synapse density after both 1 hr ($p < 0.0167$) and 24 hr ($p < 0.0167$) upscaling, and no significant difference between 1 hr and 24 hr upscaling (*$p<0.0167$; **$p<0.0033$; ***$p<0.00033$). The same description applies for all following box plots unless otherwise noted. **(B)** Column scatter plots of synapse density from all dendritic branches in all cell replicates (each column corresponding to the dendritic branches from a cell replicate). **(C)** Graphs showing all the dendritic branches (labelled as D1…D9) and corresponding synapses (superimposed colored circles along the dendrites) identified. Black hollow circles indicate excluded synapses and branches: PSD puncta more than 2 μm away from the dendrites (as identified by MAP2 immunostaining) were excluded; out-of-focus dendritic branches were excluded.

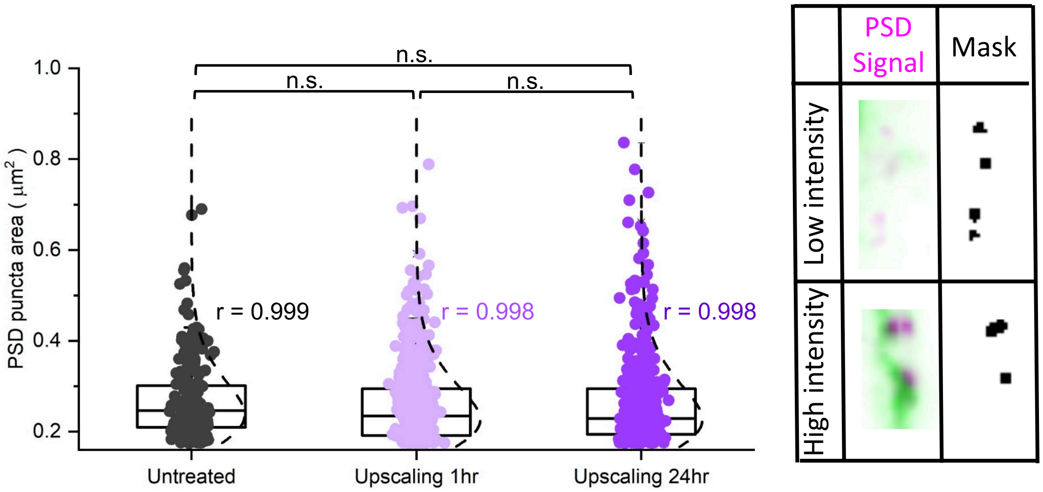


**Fig. S4.**

**Characterization of PSD95 puncta size.** Scatter plots indicating the measured PSD puncta size distribution for the synapses belonging to the dendrites in untreated (black, 225 puncta) neurons, and neurons in the upscaling 1 hr (lavender, 555 puncta), and upscaling 24 hr (purple, 428 puncta) conditions. There was no significant difference between the groups (Unpaired T-tests corrected for multiple testing with Bonferroni adjustment). r values indicate goodness-of-fit to a log-normal distribution (indicated by dashed lines). Right inset show examples of diffraction-limited PSD95 puncta and their corresponding identifications by local thresholding, regardless of local intensity differences.

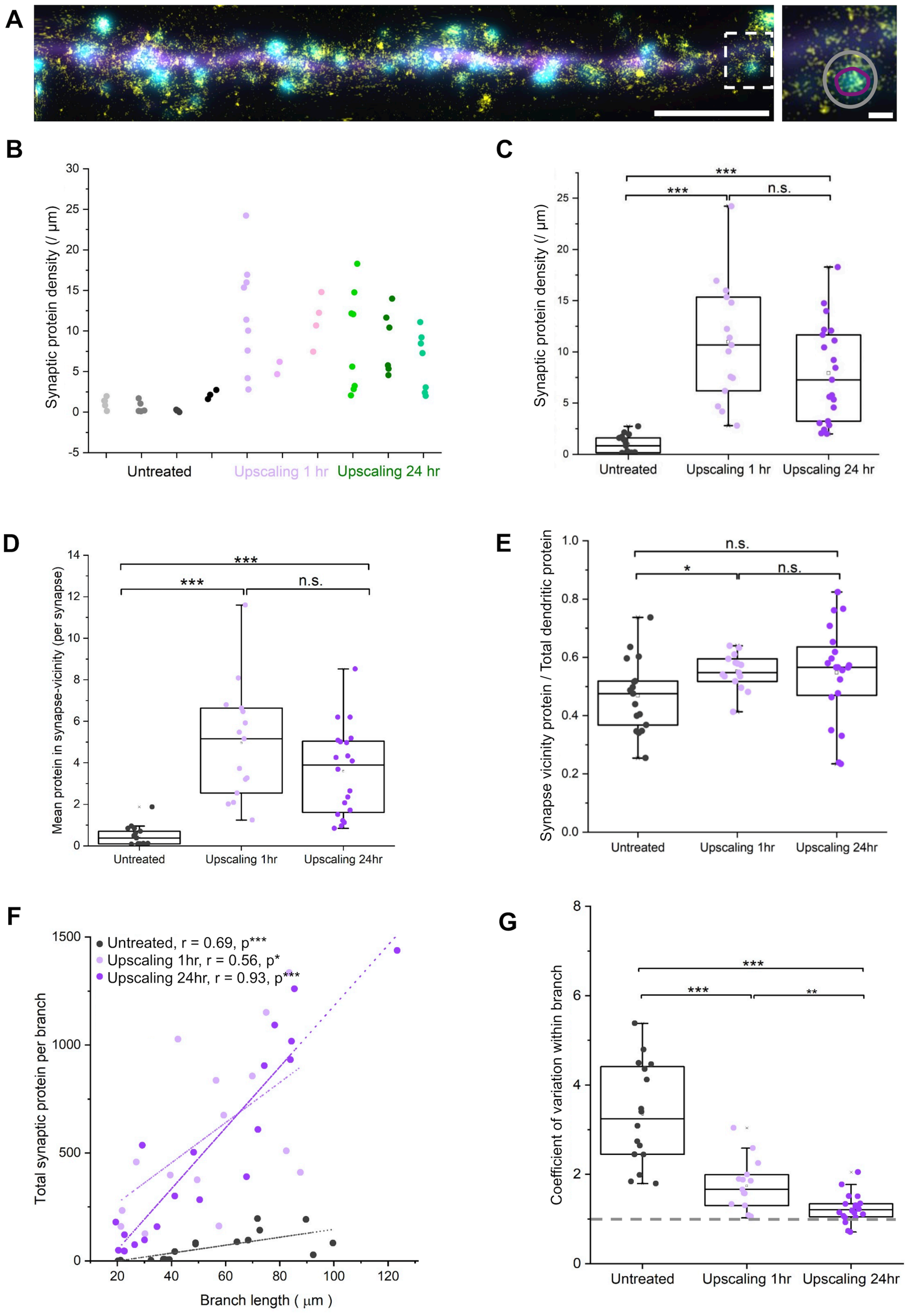


**Fig. S5.**

**Dendrite-level analysis of synaptic proteins. (A)** An example straightened dendritic branch with all its protein localizations (magenta: MAP2 immunolabel; cyan: PSD95 immunolabel; yellow: all linked protein localizations without cluster identification). Total dendritic protein localizations include localizations that are within 2.5 µm from the center line of branch (see methods). Scale bar: 5 µm. The white dashed box indicates the region corresponding to the right inset. Right inset: localizations in the synapse (purple circle) and in an expanded synapse-vicinity (grey circle). Scale bar: 0.5 µm. **(B)** Column scatter plots of synaptic protein densities from all dendritic branches in all cell replicates (each column corresponding to the dendritic branches of a cell replicate). **(C)** Column scatter plots of synaptic protein density per unit length. There were significant increases after both 1 hr ($p<0.00033$) and 24 hr ($p<0.00033$) upscaling (Unpaired T-test corrected for multiple comparisons; *$p<0.0167$; **$p<0.0033$; ***$p<0.00033$; same applies for following data). **(D)** Column scatter plots of protein in the synapse-vicinity (see main text and method) per synapse for 17 dendritic branches from 4 untreated cells (grey), 15 branches from 3 cells treated with TTX for 1 hr (light magenta), 21 branches from 3 cells treated with TTX. There were significant increases after both 1 hr ($p <0.00033$) and 24 hr ($p <0.00033$) upscaling (Unpaired T-test corrected for multiple comparison). **(E)** Column scatter plots of synaptic vicinity fractions for 17 dendritic branches from 4 untreated cells (grey), 15 branches from 3 cells treated with TTX for 1 hr (light magenta), 21 branches from 3 cells treated with TTX. There was a significant increase after 1 hr upscaling ($p <0.05$) (Unpaired T-test corrected for multiple comparisons). **(F)** Scatter plots showing significant correlations between synaptic protein of each branch (‘synaptic protein’, Y) and the branch length (µm) (X) with corresponding adjusted Pearson’s correlation coefficients r and p value ranges from one-tail tests. * and *** correspond to $p <0.05$ and $p <0.0001$, respectively. **(G)** Scatter plots indicating the coefficient of variation (CV; standard deviation divided by the mean value) of synaptic protein allocation within dendritic branches. Grey dash line indicates CV = 1. There were significant decreases after both 1 hr ($p <0.00033$) and 24 hr ($p <0.00033$) upscaling (Unpaired T-test corrected for multiple comparisons).

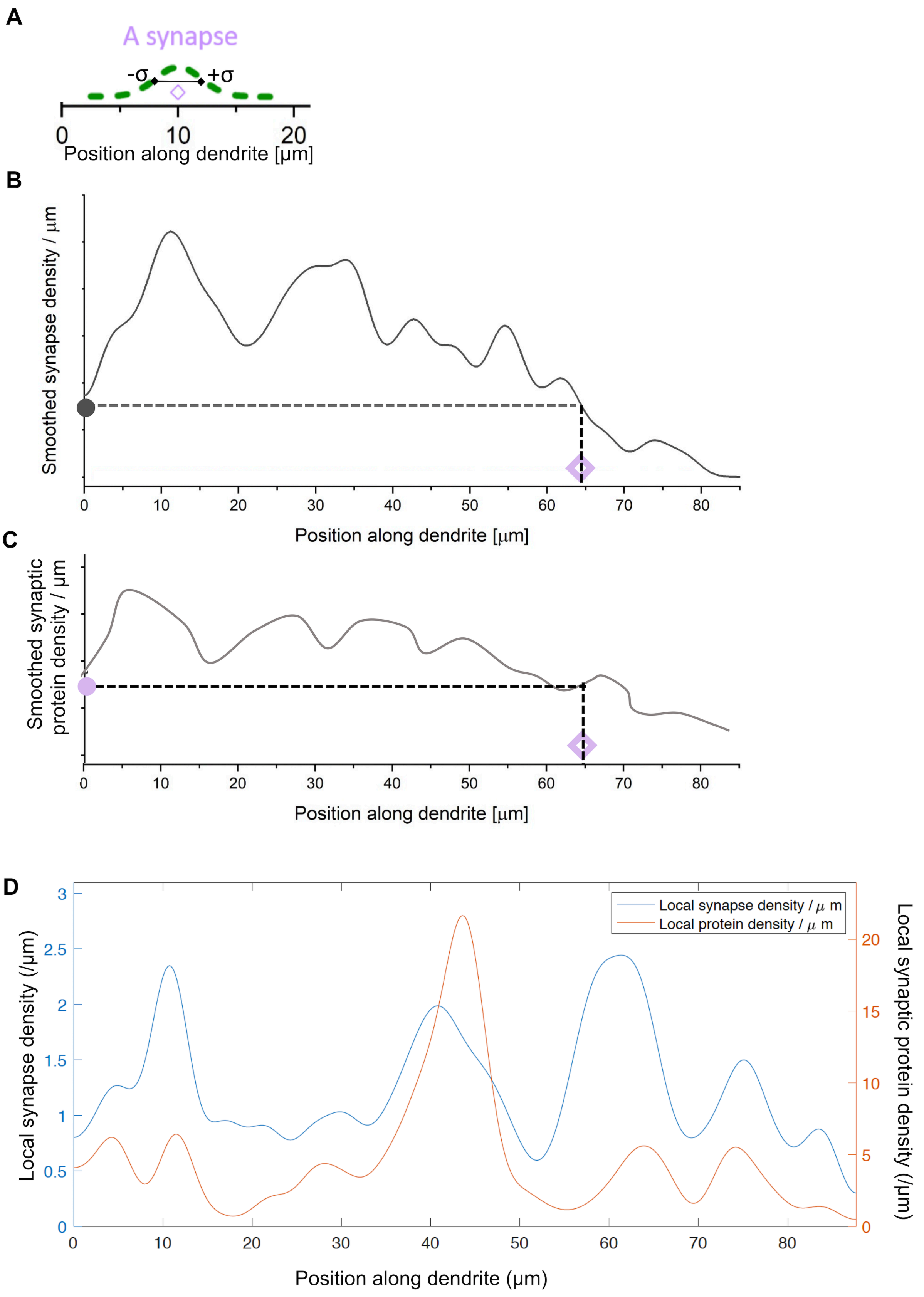
A
A synapse
-σ
+σ
0
10
20
Position along dendrite [µm]
B
Smoothed synapse density / µm
0
10
20
30
40
50
60
70
80
Position along dendrite [µm]
C
Smoothed synaptic protein density / µm
0
10
20
30
40
50
60
70
80
Position along dendrite [µm]
D
Local synapse density (/µm)
Local synaptic protein density (/µm)
Local synapse density / µ m
Local protein density / µ m
0
0.5
1
1.5
2
2.5
3
0
5
10
15
20
0
10
20
30
40
50
60
70
80
Position along dendrite (µm)

**Fig. S6.**

**Density fluctuations of synapse and synaptic protein density. (A)** Every synapse's contribution to synapse density is approximated as a unit Gaussian with a standard deviation, $\sigma$ = 2 μm. X axis indicates the position of a synapse along the dendrite. **(B)** The fluctuation of synapse density (Y axis) along a dendritic branch (X axis) generated from the superposition of all synapse Gaussians. Knowing a synapse's location on a dendrite (e.g. hollow pink diamond), one can measure the local synapse density (black dot on the Y axis). **(C)** The fluctuation of synaptic protein density (Y axis) along a dendritic branch (X axis) generated from the superposition of all synapse Gaussians using a similar method described in B, with each synapse's contribution proportional to its protein allocation level. Knowing a synapse's location on a dendrite (e.g. hollow pink diamond), one can measure the local synaptic protein density (pink dot on the Y axis). **(D)** Fluctuations of synapse density and synaptic protein density along an example dendritic branch from the 1 hr upscaling group. Notice the lack of general increase of synaptic protein density with increasing proximity-to-the-soma (towards position 0 on x axis), consistent with Fig. S2C.

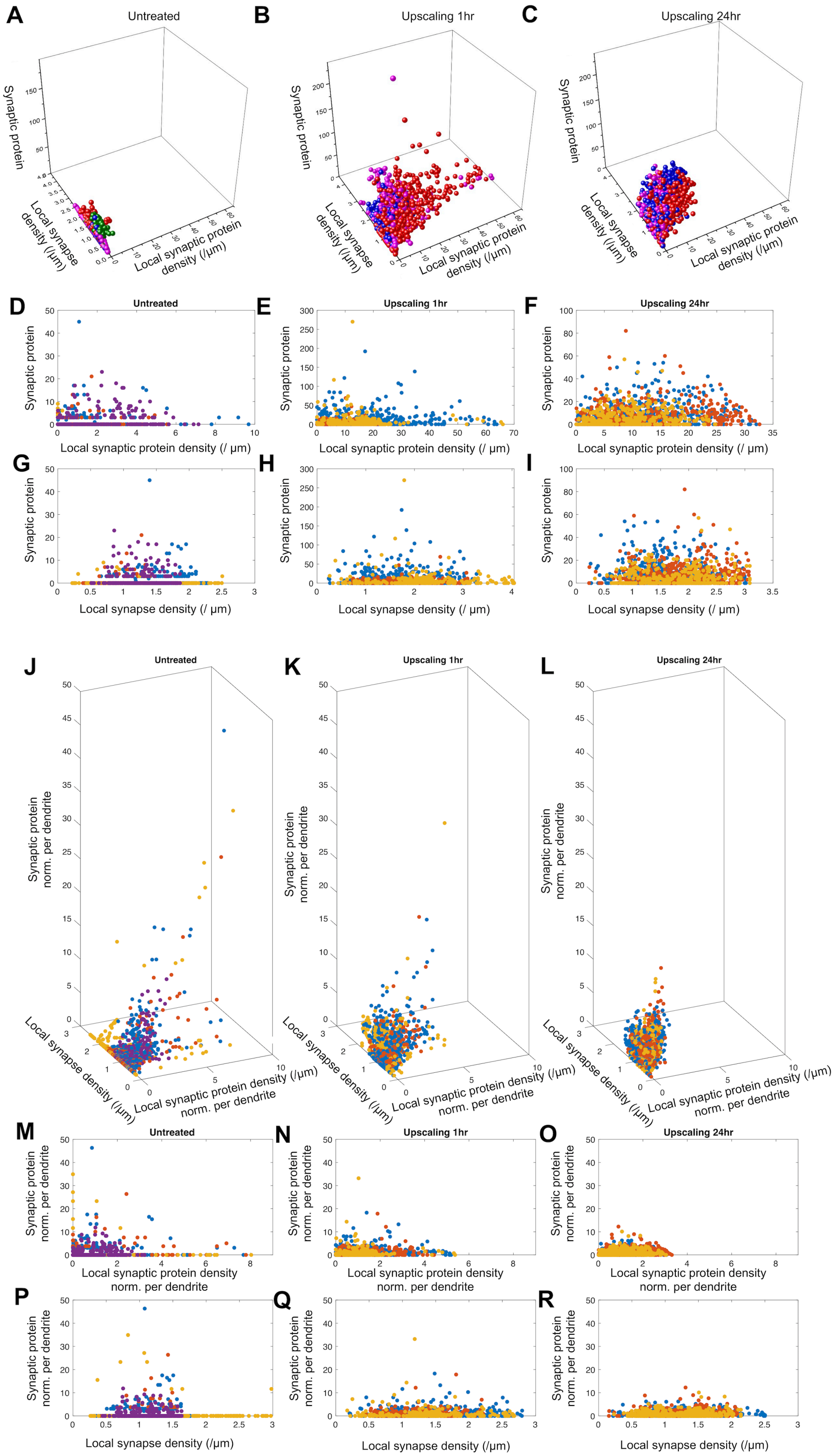
A
Untreated
B
Upscaling 1hr
C
Upscaling 24hr
Synaptic protein
Local synapse density (/µm)
Local synaptic protein density (/µm)
D
E
F
Local synaptic protein density (/ µm)
G
H
I
Local synapse density (/ µm)
J
K
L
Synaptic protein norm. per dendrite
Local synaptic protein density (/µm) norm. per dendrite
M
N
O
Local synaptic protein density norm. per dendrite
P
Q
R
Local synapse density (/µm)

**Fig. S7.**

**Synapse-level analysis of synaptic protein allocation. (A)-(C)** 3D scatter plots of **(A)** 1000 synapses from 4 untreated neurons (synapses from each neuron share the same color); **(B)** 1037 synapse from 3 upscaling 1 hr neurons; and **(C)** 1414 synapses from 3 upscaling 24 hr neurons. X axis: Local synaptic protein density; Y axis: Local synapse density; Z axis: synaptic protein of an individual synapse. **(D)-(F)** shows the scatter plots of synaptic protein vs synaptic protein density for the untreated (D), upscaling 1 hr (E), and upscaling 24 hr (F) conditions. Each color represents a cell replicate. **(G)-(I)** shows the scatter plots of synaptic protein vs synapse for the untreated (G), upscaling 1 hr (H), and upscaling 24 hr (I) conditions. Each color represents a cell replicate. **(J)-(L)** 3D scatter plots of **J)** 1000 synapses from 4 untreated neurons (synapses from each neuron share the same color); **K)** 1037 synapse from 3 upscaling 1 hr neurons; and **L)** 1414 synapses from 3 upscaling 24 hr neurons. X axis: Local synaptic protein density normalized per dendrite; Y axis: Local synapse density; Z axis: synaptic protein of a synapse normalized per dendrite. **(M)-(O)** shows the scatter plots of normalized synaptic protein vs normalized local synaptic protein density for the untreated (M), upscaling 1 hr (N), and upscaling 24 hr (O) conditions. Each color represents a cell replicate. **(P)-(R)** shows the scatter plots of normalized synaptic protein vs local synapse density for untreated (P), upscaling 1 hr (Q), and upscaling 24 hr (R). Each color represents a cell replicate.

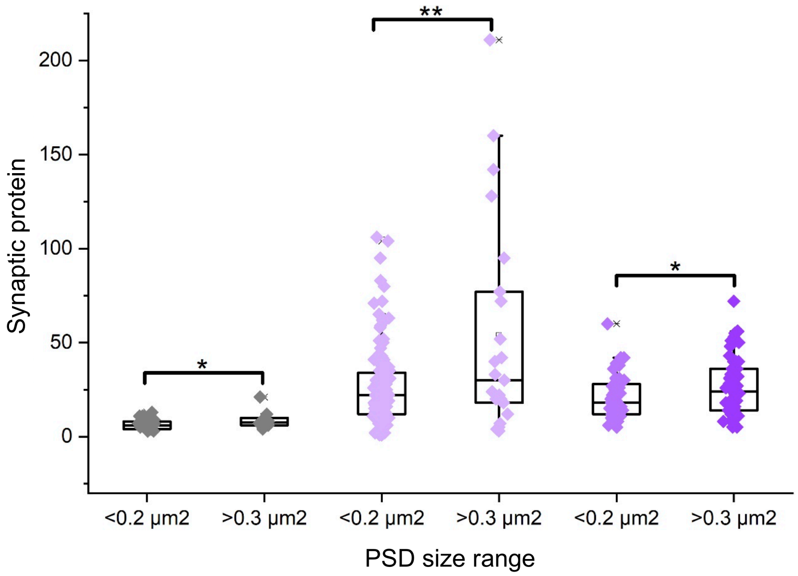


**Fig. S8.**

**Analysis of synapse-size influence on synaptic protein allocation.** Column scatter plots showing all the non-zero synaptic protein allocations of synapse with smaller (< 0.2 $\mu m^2$) and larger (>0.3 $\mu m^2$) measured using PSD-95 area (proxy for spine volume- see Methods) in the untreated (black), 1hr upscaling (light purple), and 24hr upscaling (dark purple) conditions. Unpaired two-sample T-tests were performed within each treatment group. * and ** indicate $p < 0.05$ and $p < 0.01$, respectively.

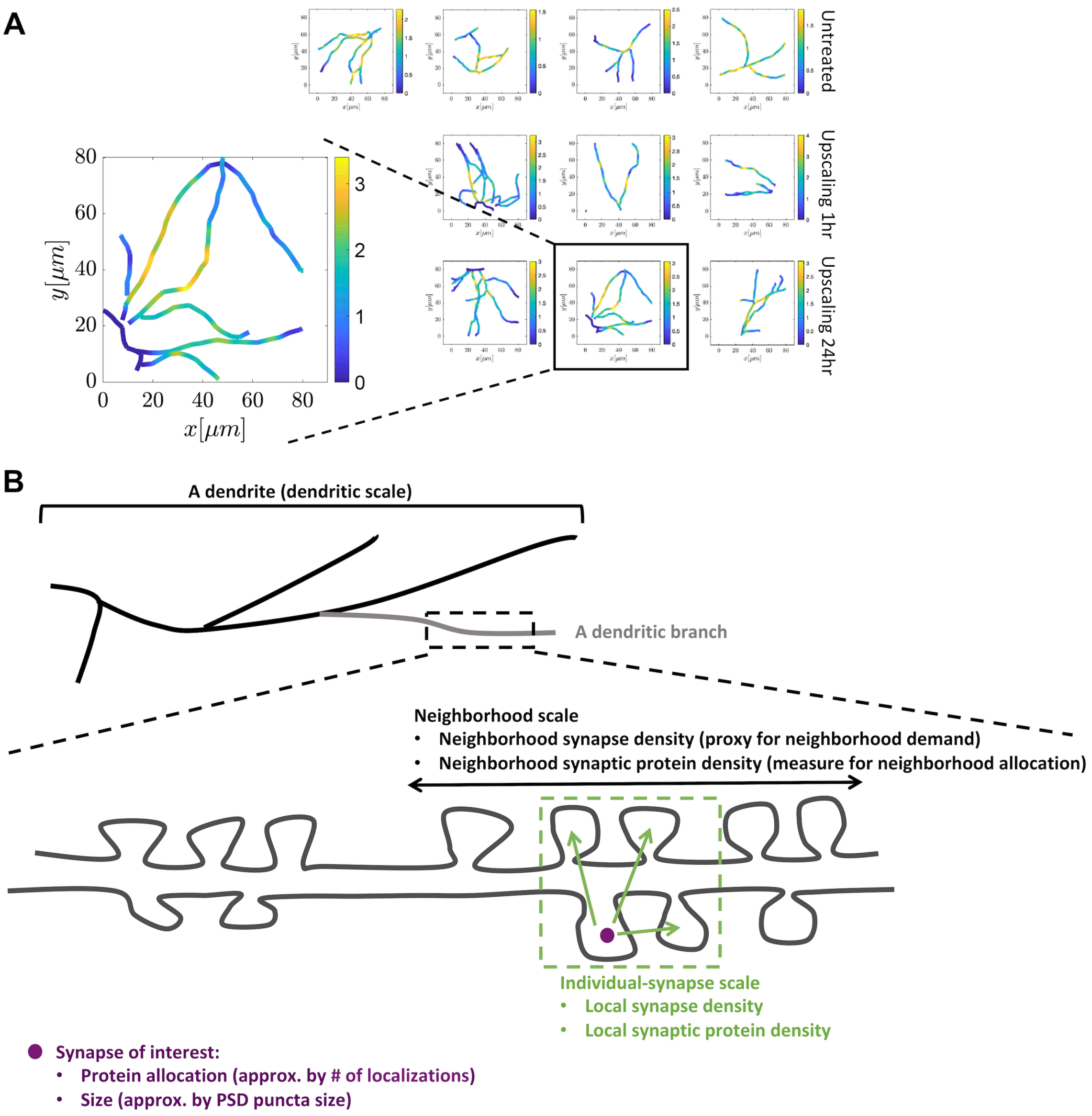


**Fig. S9.**

**Multi-scale analysis of synaptic-protein allocation. (A)** Heatmaps of synapse density (per µm dendrite; color scale unit: per µm; example dendrite corresponding to Fig. 4A was enlarged at left) exhibit spatial heterogeneity at intermediate length-scales ($10^0$ - $10^1$ µm). (B) Illustration of the dendrite scale, the neighborhood scale, and the individual synapse-scale (the purple dot indicates a synapse-of-interest and its local neighbors).

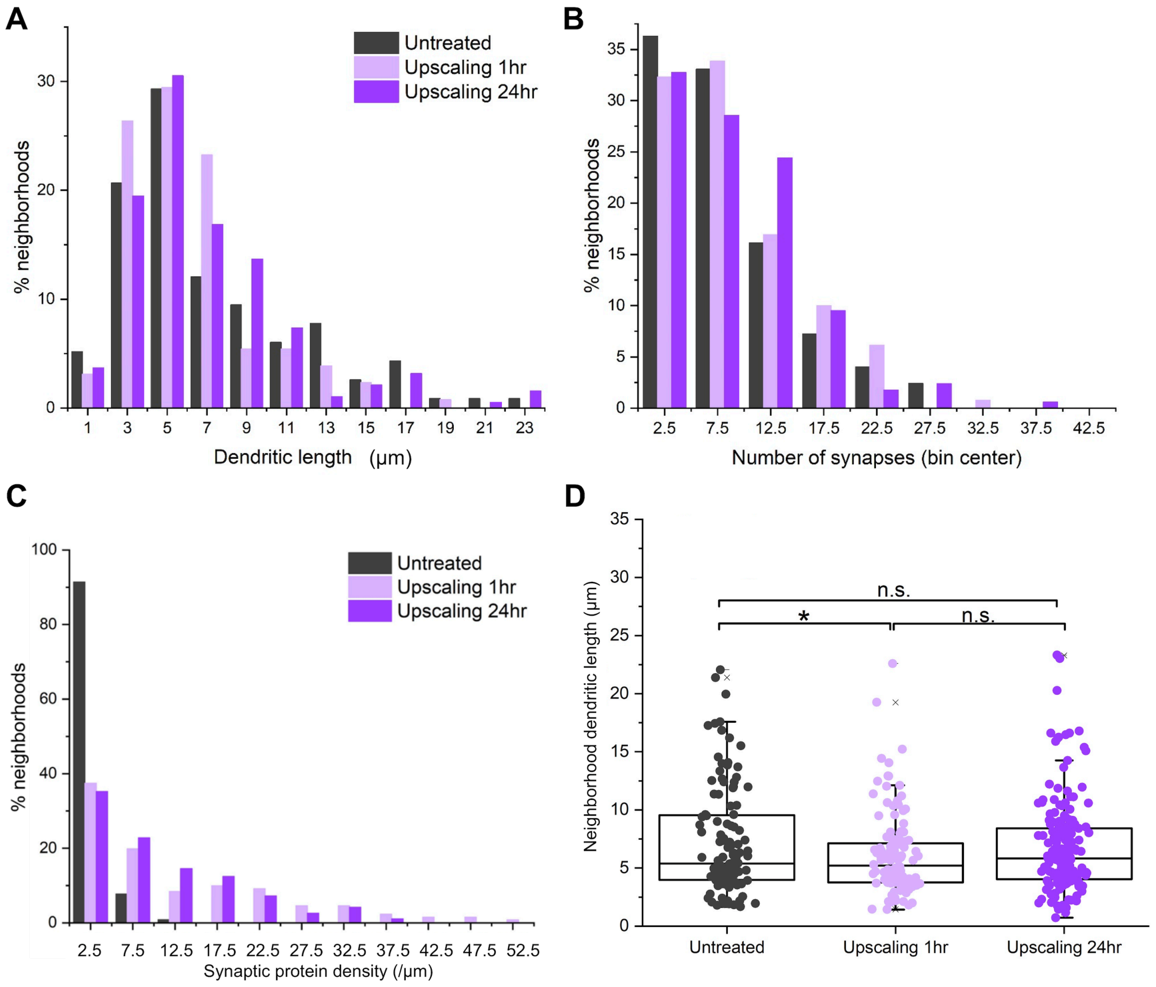


**Fig. S10.**

**Characterization of synapse neighborhoods. (A)** Histogram showing the fraction of neighborhoods that are the indicated by the dendritic length in each condition. Mean length = 7.2 µm for control; mean length = 6.2 µm for 1 hr upscaling; mean length = 6.8 µm for 24 hr upscaling. **(B)** Histogram showing the fraction of neighborhoods that possesses the indicated number of synapses in each condition. Mean = 7.9 for control; mean = 7.9 for 1 hr upscaling; mean = 7.35 for 24 hr upscaling. **(C)** Histogram showing the fraction of neighborhoods that possesses the indicated neighborhood synaptic protein distributions. Mean = 2.4 for control; mean = 12.0 for 1 hr upscaling; mean = 9.2 for 24 hr upscaling. 1 hr upscaling distribution stretched into the high-density region, introducing higher variability. **(D)** Scatter plots showing the mean dendritic lengths of neighborhoods for untreated (grey), 1 hr upscaling (lavender), and 24 hr upscaling (purple). There was a significant decrease of neighborhood size (by dendritic length) following 1 hr upscaling ($p<0.0167$) compared to control, but not after 24 hr upscaling (Unpaired T-test corrected for multiple comparison).

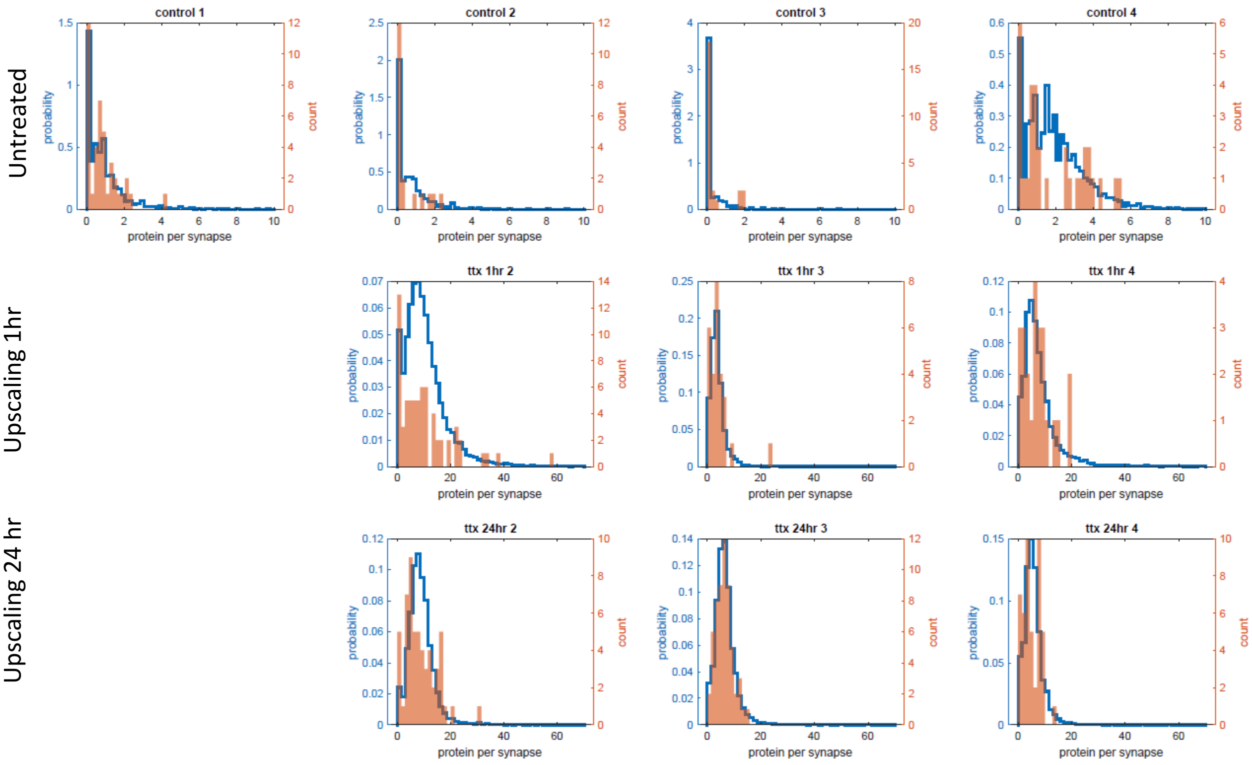


**Fig. S11.**

**Randomization of synaptic protein allocation.** Histograms showing the distributions of mean neighborhood synaptic protein (per synapse) for 500 randomized cases (blue lines) and in the original dendrites (orange bars) for each replicate in the control, upscaling 1 hr and upscaling 24 hr conditions.

**Table S1.**

Coefficients of variation (CV) for neighborhoods' synaptic-protein-per-synapse

| | Untreated | Upscaling 1hr | Upscaling 24 hr |
|---|---|---|---|
| CV of neighborhood synaptic-protein per-synapse | 1.31 | 1.04 | 0.68 |
